# Seed size and source reduction during seed filling effect on quality traits of winter and spring rapeseed

**DOI:** 10.64898/2026.02.21.707178

**Authors:** José Verdejo, Daniel F. Calderini

## Abstract

**CONTEXT:** Rapeseed is a globally significant oil crop, exhibiting highly plastic responses among seed yield components (seed number and weight). However, there remains a notable gap in knowing the distribution of quality traits among seed size categories and understanding how seed size and source-sink (S-S) ratio influence comprehensive seed quality traits.

**OBJECTIVE:** This study investigated the effects of seed size and S-S ratio reduction on the quality traits of winter and spring rapeseed genotypes.

**METHODS:** The experiments were carried out at field conditions in Valdivia, Chile, where seed yield, yield components, oil, protein, and element concentrations (P, K, S, Ca, Mg, B, Cu, Fe, Mn, Zn, and Na) were evaluated across five seed size categories; very small (< 1.4 mm), small (1.4–1.7 mm), medium (1.7–2.0 mm), large (2.0–2.36 mm), and very large (> 2.36 mm). Treatments included a control and a reduced S-S ratio (75% shading), which significantly increased seed weight (P < 0.05).

**RESULTS:** Both genotype and seed size affected (P< 0.050) the quality traits. Larger seeds exhibited higher Mg and B concentrations, as well as lower K, Ca, Fe and Na. Shading affected seed size distribution, favouring a higher proportion of large seeds. Under the shading treatment, the small seed category reached 5% lower oil concentration, while protein seed concentration increases 6% in both genotypes. Principal component analysis highlighted the complex interaction between yield, yield components, and quality traits, since there was no clear separation between different seed size categories and S-S ratio treatments.

**CONCLUSION:** These results provide insights into the plasticity of rapeseed quality traits, highlighting their collective impact on nutrient profiles.

**SIGNIFICANCE:** This information is helpful for optimising cultivation practices and informing breeding programmes aimed at improving seed quality, particularly in high-yielding environments susceptible to environmental stresses.

**Graphical abstract:** 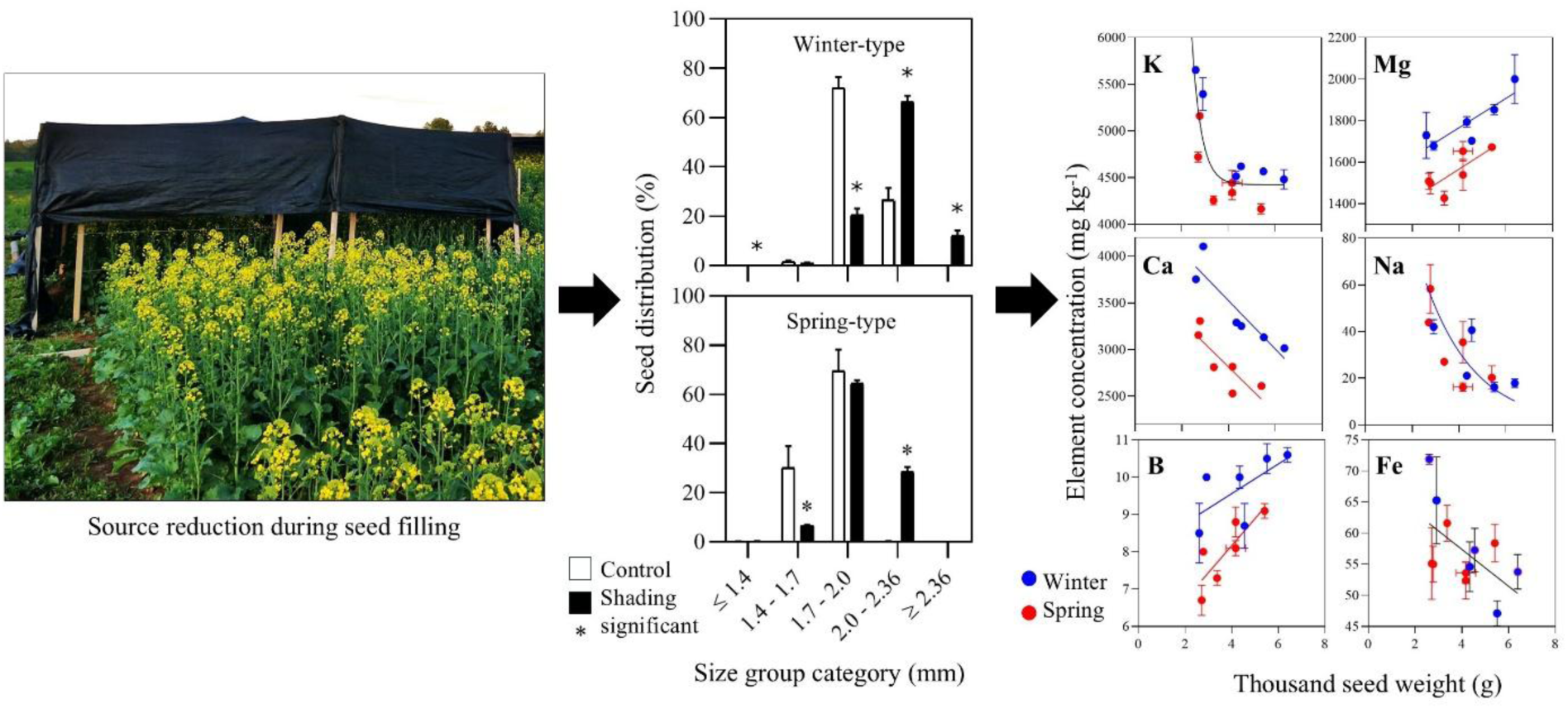

## 1. Introduction

Rapeseed is an important global oil crop with several advantages, as it offers the highest unsaturated fatty acid ratio among plant oils (Farsak, 2009). Rapeseed oil contains approximately 65% oleic, 20% linoleic, 9% linolenic, and 2% stearic acids (Weiss, 2000), making it nutritionally rich for human consumption (Beyzi et al., 2019). The meal obtained after oil extraction contains a protein concentration of 36–39% (Newkirk, 2011), making it a viable and economical protein supplement for replacing soybean in formulated animal feeds (Vahedifar and Wu, 2022). Additionally, rapeseed is used as an industrial oil, with expanding markets driving the development of fatty acid profiles suitable for biodiesel and industrial feedstocks (Fobert et al., 2008; Thiyam-Hollaender et al., 2013).

Rapeseed is a very plastic crop showing compensations between seed yield components. For instance, full compensation by seed weight was reported by Labra et al. (2017) when seed number decreased in response to shading in Southern Chile. Partial compensation was also reported in this crop by other authors (Kirkegaard et al., 2018; Verdejo and Calderini, 2020). However, it remains largely unknown how this plasticity impacts the comprehensive seed quality traits, including not only oil and protein but also macro- and micronutrient concentrations. This knowledge gap is particularly critical given the increasing prevalence of environmental changes, such as reductions in solar radiation observed in high-yielding regions like Southern Chile (Rivelli et al., 2021), which directly influence resource availability during seed filling.

Oil seed concentration of rapeseed was reported negatively affected by abiotic constraints such as increased temperatures or reduced radiation (Triboi-Blondel and Renard, 1999; Kirkegaard et al., 2018; Pokharel et al., 2021). When this quality trait was compromised, protein seed concentration responded positively, demonstrating a trade-off between both numerical yield components (Rondanini et al., 2014; Labra et al., 2017; Kirkegaard et al., 2018). For example, when the temperature was increased by 8 °C, a decrease in oil concentration in the seeds was found, ranging from 3 to 6 percentage points, with a concurrent increase in seed protein concentration of 4–7 percentage points (Triboi-Blondel and Renard, 1999; Pokharel et al., 2021). Additionally, Kirkegaard et al. (2018) reported a 40–50% decrease in seed oil concentration and an increased protein concentration by 30–40%, relative to the control treatment, in experiments where the source-sink ratio (S-S ratio) was reduced by 85% during the critical period of seed number determination (100-500 °Cd after the start of flowering).

These findings, however, contrast with data from the experiment carried out by Labra et al. (2017) in Southern Chile, where high stability of seed quality traits (seed oil and protein concentrations) was reported when the S-S ratio was decreased at the beginning of seed filling. The conservative response of seed oil concentration under different S-S ratios was also reported for sunflower by Castillo et al. (2017) in Southern Chile, a region recognised as one of the most favourable areas for temperate grain crops globally (Sandaña et al., 2009; Quintero et al., 2018; Verdejo and Calderini, 2020; del Pozo et al., 2022). In this region, rapeseed is a highly adapted and sown crop (Mera et al., 2015). Therefore, the sensitivity of both oil and protein seed concentrations in response to the environment is still under discussion.

While the majority of rapeseed research has predominantly focused on oil and protein content, a holistic assessment of seed quality, essential for global food and feed security (Hefferon, 2015; Kreitzman et al., 2020; FAO et al., 2023), mandates the inclusion of the full elemental composition of seeds (e.g., phosphorus, potassium, sulphur, calcium, magnesium, iron, manganese, zinc and copper). Moreover, seed size and nutrients concentration are important for seedling establishment and growth (Zhang et al., 2007), regarding the requirement of both farmer and industrial markets (Richards and Lukacs, 2002). Larger seeds have been proposed as an effective breeding strategy to increase seed yield of oil crops (López Pereira et al., 1999; Zhu et al., 2011). Furthermore, Beyzi et al. (2019) reported that larger seeds of rapeseed had higher concentration of calcium, magnesium, potassium, and sulphur than smaller ones. Similar results were found in wheat (Calderini and Ortiz-Monasterio, 2003; Larroque et al., 2022). In contrast, small seeds have lower emergence capacity and reduced agricultural performance (Gulden et al., 2004; Lamb and Johnson, 2004), negatively affecting animal digestibility (Jensen et al., 1995) and produce oil that deteriorates quickly due to oxidation and hydrolysis processes (Rotkiewicz et al., 2002). This because of the relatively small embryos in small seeds with lower reserve compounds (Mińkowski, 2000; Batool et al., 2022). Seed size information is of high interest since Panthee et al. (2005) demonstrated that this trait impacts on oil and protein concentrations in soybean, showing a positive correlation between protein concentration and seed size, which in turn affects seed oil concentration. However, partial information is available in this regard. For example, Beyzi et al. (2019) evaluated only one rapeseed cultivar considering seed quality traits (oil, protein, and element concentrations) across different seed size categories, while Kowalska et al. (2020) carried out a seed quality appraisal (protein and element concentrations) of fifteen winter-type genotypes omitting the consideration of seed oil concentration and seed size categories. None have comprehensively addressed the interactive effects of seed size categories and source-sink ratio manipulations on the complete spectrum of oil, protein, and elemental concentrations across diverse winter and spring rapeseed genotypes.

As it was pointed out above, seed weight and size of rapeseed is plastic in response to seed number variation (Fig. 3 in Labra et al., 2017; Rivelli et al., 2024). Therefore, any biotic or abiotic constrain during the critical period for seed number determination of rapeseed (Kirkegaard et al., 2018), could affect not only seed yield but also seed size, weight and its quality traits. Verdejo and Calderini (2020) reported different periods for seed weight plasticity in response to the S-S ratio for winter and spring genotypes, specifically 15–30 and 0–15 days after flowering (DAF), respectively. In the winter-type genotypes, seed weight increased by 28% to 33% over the control under the 15–30 DAF shading treatment, whereas in spring types, seed weight rose from 15% to 39% during the 0–15 DAF treatment. Furthermore, they reported resilience of seed yield during these seed filling periods, due to the ability of seed weight to increase in response to reductions of seed number under lower source-sink ratio. However, little is known about the sensitivity of seed size, and the concentration of oil, protein and elements (P, K, S, Ca, Mg, B, Cu, Fe, Mn, Zn, and Na), when seed weight is increased under seed number reduction in rapeseed. Therefore, this study directly addresses this knowledge gap by comprehensively investigating whether the well-documented increase in seed weight due to source-sink ratio reductions (e.g., shading) modifies the oil, protein, and element concentrations across different seed size categories in both winter and spring rapeseed genotypes. By doing so, we aim to uncover the underlying nutritional plasticity of rapeseed and its implications for optimising cultivation practices and breeding programmes under changing environmental conditions.

## 2. Materials and Methods

### 2.1. Experimental set-up and plant materials

In the present study, the assessed seeds of rapeseed were harvested in previous experiments evaluating the impact of the decreased S-S ratio after flowering on winter and spring genotypes (Verdejo and Calderini, 2020). Briefly, the field experiments were carried out at the Austral Experimental Station in Valdivia, Chile (39° 47’ S, 73° 14’ W). Two oilseed genotypes were evaluated in this study, i.e. Mercedes (winter-type) and Solar CL (spring-type). The winter genotype was sown on May 9^th^, 2018 in plots measuring 2 m in length and 3.5 m in width, with 7 rows spaced 0.3 m apart and sown at a density of 35 plants m^-2^. The spring genotype was sown on September 3^rd^, 2018, in similar plots than Mercedes, but with 11 rows spaced 0.175 m apart (seed rate of 55 plants m^-2^). One month before the experiments, 2 t ha^-1^ of CaCO_3_ was applied in the soil to prevent low pH (<6.2). At sowing, 150 kg ha^−1^ of Ca(H_2_PO_4_)_2_ was fertilized in each experiment to prevent nutritional phosphorus shortage. Nitrogen was applied twice during the crop cycle, i.e. 100 kg N ha^−1^ at the plant emergence stage (BBCH 10) and again, 100 kg N ha^−1^ more were fertilized when the fifth internode was expanded (BBCH 35). Nitrogen application in both experiments was in the form of (NH_4_)NO_3_+ CaCO_3_+ MgCO_3_ (50% NO_3_ and 50% NH_4_). No additional nutrients were added, as the soil had sufficient levels according with the soil analysis. Plots were surface irrigated until physiological maturity with a drip tape system as needed, depending on rainfall, to prevent water shortage. Optimal management practices were implemented to prevent or control diseases, pests, and weeds in accordance with the rates recommended by the manufacturers, ensuring the crops remained free of biotic constraints.

Treatments consisted of a control and a reduction of the S–S ratio, set after the start of flowering, specifically 0–15 and 15–30 DAF (days after flowering). The experiments were carried out using a randomized block design with three replicates. The S-S ratio was reduced by shading the plots with black nets that intercepted 75% of solar radiation reaching the top of the crop. The black nets were set at 0.3-0.4 m above the top of the plants during the treatment, and the south side was left open to allow air circulation and pollinator access, as described by Labra et al. (2017). The treatments evaluated in this study were the control and the S–S ratio reduction during the most effective period for the increase of seed weight, i.e. 15–30 DAF in the winter genotype Mercedes and 0–15 DAF in the spring genotype Solar CL (Verdejo and Calderini, 2020). Seed samples were harvested from one linear metre of the central rows of each plot at maturity (BBCH 89), when the seeds inside the siliques were dark and hard. The seed number was measured after oven-drying the samples at 65 °C for 48 hours using a seed counter (Pfeuffer GmbH, Kitzengen, Germany). Subsequently, seed yield was measured, and the thousand seed weight (TSW) was estimated as the ratio between seed yield and seed number. These seeds were stored at 5 °C in sealed plastic bags in a refrigerator. Seed yield, seed number, weight and other crop traits were reported in Verdejo and Calderini (2020).

### 2.1 Seed size categories and quality traits determination

Seed size was classified into five categories, i.e. very-small, small, medium, large and very-large (Table 1) by using a motorised sieve shaker (W.S. Tyler, Ohio, USA). The seed size categories from the present study were primarily derived from Beyzi et al. (2019) to facilitate comparisons with data from the literature. Quality traits, such as oil, protein, and nutrient concentrations were determined for the small, medium, and large size categories, as these sizes were shared across the genotypes and due to the very few seeds available from the extreme categories, very-small and very-large in Mercedes (winter-type), and mainly very large in Solar CL (spring-type).

**Table 1.** Seed yield, seed number and thousand seed weight in control and reduced source-sink ratio (S-S ratio) treatments of winter-type (Mercedes 15-30 DAF) and spring-type (Solar CL 0-15 DAF) rapeseed genotypes in each seed size category.

|  |  | Seed yield (g m<sup>-2</sup>) |  |  |  |  |  |
| --- | --- | --- | --- | --- | --- | --- | --- |
| Genotype | Shading | Very small<br>≤ 1.4 | Small<br>1.4 - 1.7 | Medium<br>1.7 - 2.0 | Large<br>2.0 - 2.36 | Very large<br>≥ 2.36 | Total |
| Winter-type | Control | 0.1 ± 0.1 a | 12.7 ± 3.3 a | 602.2 ± 32.9 a | 224.9 ± 52.5 b | 0.2 ± 0.1 b | 840.2 ± 48.3 a |
|  | S-S ratio | 0.4 ± 0.2 a | 4.7 ± 2.3 a | 125.8 ± 15.6 b | 410.1 ± 45.5 a | 78.4 ± 22.0 a | 619.4 ± 71.1 a |
| Spring-type | Control | 0.9 ± 0.3 a | 178.7 ± 19.1 a | 472.0 ± 119.5<br>a | 1.8 ± 0.8 b | 0.0 ± 0.0 a | 653.4 ± 102.1<br>a |
|  | S-S ratio | 1.3 ± 0.6 a | 32.9 ± 4.1 b | 323.4 ± 25.0 a | 143.9 ± 16.9 a | 0.2 ± 0.1 a | 501.7 ± 42.1 a |
|  |  | Seed number (10<sup>3</sup> m<sup>-2</sup>) |  |  |  |  |  |
|  |  | Very small<br>≤ 1.4 | Small<br>1.4 - 1.7 | Medium<br>1.7 - 2.0 | Large<br>2.0 - 2.36 | Very large<br>≥ 2.36 | Total |
| Winter-type | Control | 0.1 ± 0.1 a | 4.4 ± 1.1 a | 141.0 ± 8.6 a | 41.3 ± 9.6 a | 0.0 ± 0.0 b | 186.9 ± 9.3 a |
|  | S-S ratio | 0.3 ± 0.1 a | 1.8 ± 0.8 a | 27.7 ± 3.6 b | 64.1 ± 6.2 a | 9.3 ± 2.4 a | 103.2 ± 9.5 b |
| Spring-type | Control | 0.6 ± 0.2 a | 66.3 ± 7.0 a | 140.6 ± 35.4 a | 0.4 ± 0.2 b | 0.0 ± 0.0 | 208.0 ± 29.0 a |
|  | S-S ratio | 0.7 ± 0.4 a | 11.8 ± 1.5 b | 77.5 ± 6.3 a | 26.6 ± 3.2 a | 0.0 ± 0.0 | 116.7 ± 10.3 b |
|  |  | Thousand seed weight (g) |  |  |  |  |  |
|  |  | Very small<br>≤ 1.4 | Small<br>1.4 - 1.7 | Medium<br>1.7 - 2.0 | Large<br>2.0 - 2.36 | Very large<br>≥ 2.36 | Total |
| Winter-type | Control | N/A | 2.92 ± 0.08 a | 4.34 ± 0.03 b | 5.51 ± 0.04 b | 6.90 ± 0.19 b | 4.5 ± 0.1 b |
|  | S-S ratio | 1.36 ± 0.13 | 2.60 ± 0.09 b | 4.55 ± 0.05 a | 6.39 ± 0.06 a | 8.31 ± 0.07 a | 6.0 ± 0.2 a |
| Spring-type | Control | 1.55 ± 0.13 a | 2.71 ± 0.01 a | 3.37 ± 0.01 b | 4.17 ± 0.42 b | N/A | 3.1 ± 0.1 b |
|  | S-S ratio | 1.94 ± 0.23 a | 2.78 ± 0.03 a | 4.17 ± 0.01 a | 5.41 ± 0.01 a | 6.86 ± 0.93 | 4.3 ± 0.0 a |

Mean ± standard error. Different letters indicate significant differences ( $p < 0.05$ ) between the S-S ratio treatments of each genotype. N/A = Not available due to few seeds in the seed size category.

Oil concentration was determined using Near Infrared Reflectometry (NIR) (Foss Infratec 1241, Hilleroed, Denmark), and seed nitrogen concentration was measured using the Kjeldahl method (Kirk, 1950). The protein concentration of seeds was calculated using a conversion factor of 5.8 (Merrill and Watt, 1973). Seed element concentrations (P, K, S, Ca, Mg, B, Cu, Fe, Mn, Zn, and Na) were determined using standard methods (Sadzawka et al., 2007). Concentrations of oil, protein, and elements were expressed on a dry matter basis.

### 2.3. Statistical analysis

Analysis of variance (ANOVA) and correlation analysis were conducted for seed oil, protein, and nutrient concentrations in response to the source-sink (S-S) ratio treatments for each genotype, with the aim of comparing the S-S ratio treatments and different seed size categories. Differences among the sources of variation were deemed statistically significant at a probability of 5%, by using the least squares mean differences test. Statistical analyses were performed using Statgraphics Centurion 18 (https://www.statgraphics.com/). Linear regression analyses were carried out by GraphPad Prism 8 (https://www.graphpad.com/) to evaluate the data fit, as well as the slopes and intercepts of the linear associations. The distribution normality and homogeneity of residuals across the dataset were verified through Shapiro-Wilk and Levene’s tests, respectively (Kutner et al., 2004).

Principal component analysis (PCA) was executed on a data matrix comprised of rows representing the two genotypes (Lumen and Solar CL), two S-S ratio treatments, and three seed size categories (small, medium, and large, n = 12). The columns included the observed variables for each replicate: seed yield, seed number, thousand seed weight, and seed quality traits (seed oil, protein, and element concentrations). Biplots were constructed using the first two principal components (PC1 and PC2) with JMP 11 from SAS.

## 3. Results

### 3.1 Seed yield, yield components and quality traits affected by shading

Seed crop samples used in the present study were harvested in two larger experiments, whose environmental conditions and harvest data were reported in Verdejo and Calderini (2020). To have a picture of the results reported in that article, the highlights of the key findings are briefly depicted below.

Control winter-type rapeseed (Mercedes) reached high seed yield and above-ground biomass (Fig. 1A and 1B). The 15-30 DAF S-S ratio treatment decreased (p ≤ 0.05) seed yield by 26%, but above-ground biomass was not affected (p > 0.05). Interestingly, seed yield components showed compensation as the negative effect of shading on seed number was partially compensated by the increase (p ≤ 0.05) of TSW (Fig. 1C and 1D). Specifically, seed number decreased 45%, but in contrast, TSW increased 33%. The quality traits, such as seed oil concentration (Fig. 1E and 1F), were not affected by the S-S ratio while, shading increased seed protein concentration by 1.8 percentage points over the control treatment (17.5%).

**Figure 1.**
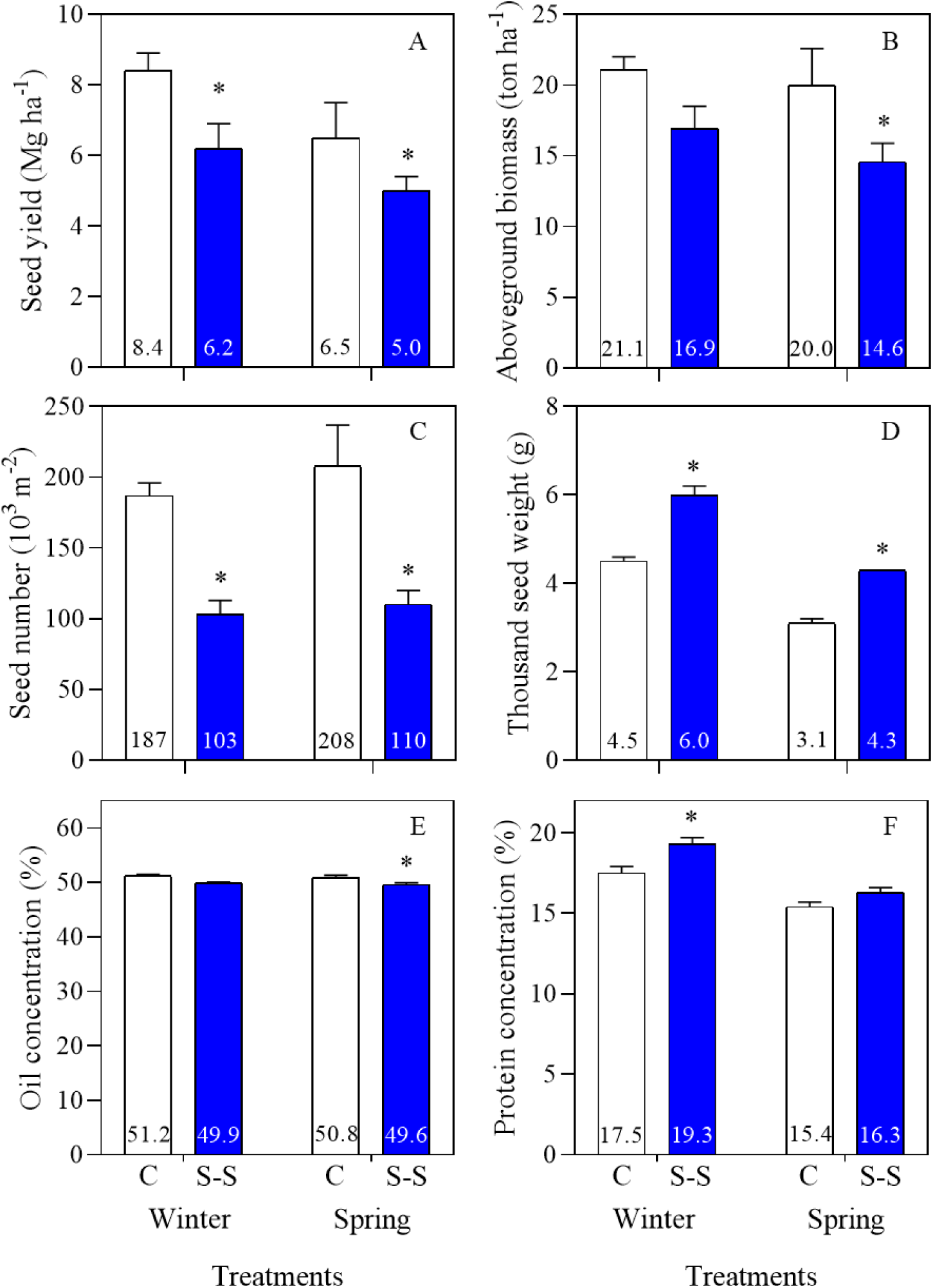
Seed yield (A), above-ground biomass (B), seed number (C), thousand seed weight (D), seed oil (E) and protein (F) concentrations of winter and spring genotypes under control (white bars) and reduced source-sink (S-S) ratio (blue bars) treatments. Asterisks indicate significant difference (p < 0.05) between the control and decreased source-sink (S-S) ratio treatments within each genotype. The segments represent the standard error of the means.

The control treatment of the spring-type rapeseed (Solar CL) achieved 6.5 Mg ha^-1^ of seed yield and high above-ground biomass (Fig. 1A and 1B). The reduction of the S-S ratio in the 0-15 DAF treatment decreased (p ≤ 0.05) seed yield and above-ground biomass by 23 and 27%, respectively. As seed yield, the number of seeds was negatively affected by shading but to a higher extent than in the winter-type, regarding that this yield component decreased 44%. On the other hand, TSW increased 39% (Fig. 1C and 1D). Additionally, shading affected seed oil concentration, which decreased 1.2 percentage points from the control (50.8%), but seed protein concentration was not modified (p > 0.05) by the S-S ratio treatment (Fig. 1E and 1F).

### 3.2 Seed yield and components across the seed size categories in the winter and spring-type genotypes affected by the S-S ratio

To have an insight on the effect of the S-S ratio treatments, seeds of each rapeseed type were divided in five seed size categories: (i) very small (< 1.4 mm), (ii) small (between 1.4 to 1.7 mm), (iii) medium (between 1.7 to 2.0 mm), (iv) large (between 2.0 to 2.36 mm) and (v) very large (>2.36 mm). The extreme categories (very-small and very-large) contributed by 6.4% to the total seed yield in the winter and 0.2% in the spring-type genotypes, averaged across both control and shading treatments (Fig. 2). Consequently, these categories were excluded from the determination of quality traits due to the limited number of seeds obtained per replication, supported by their small contribution to seed yield.

**Figure 2.**
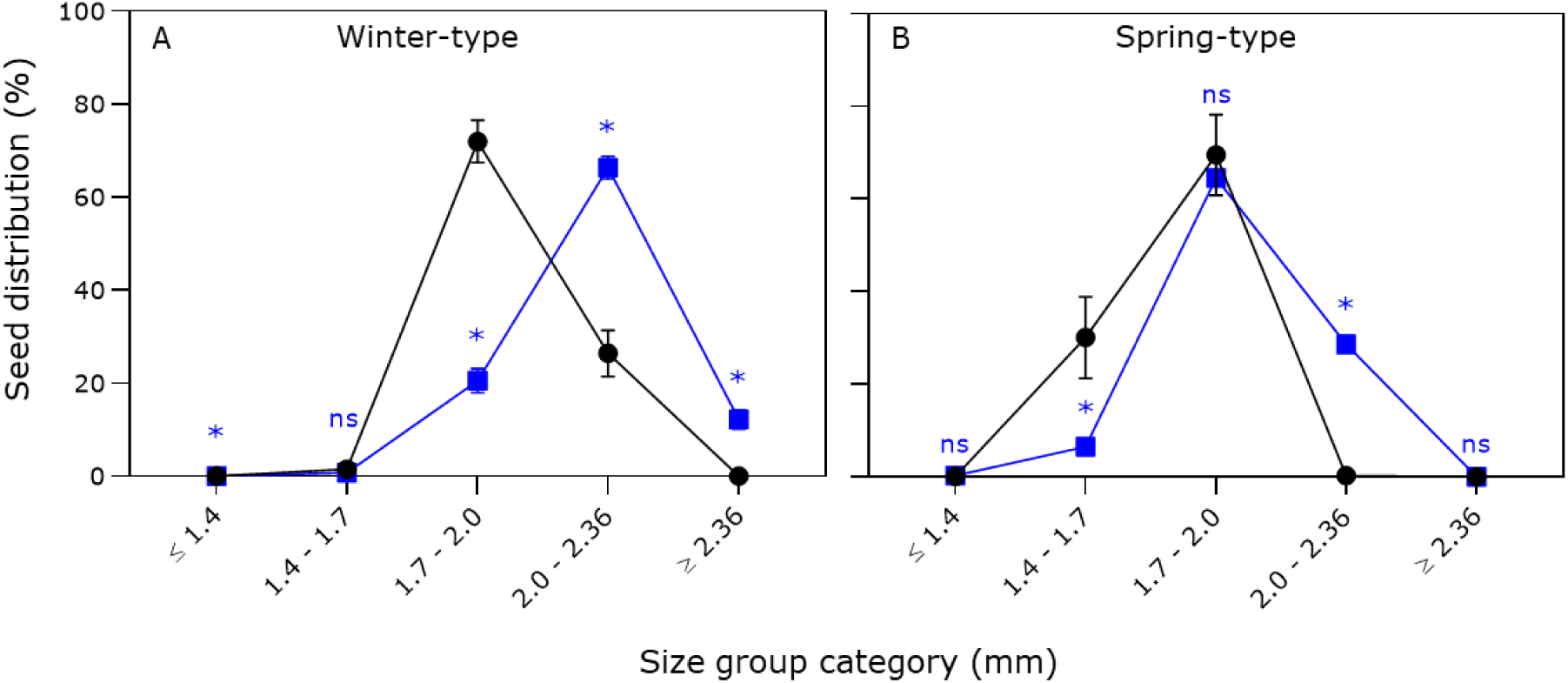
Seed distribution of winter (A) and spring (B) genotypes across the seed size categories of control (black circles) and decreased source-sink (S-S) ratio (blue squares) treatments. Asterisks indicate significant difference (p < 0.05) between the decreased source-sink (S-S) ratio and control treatments within each seed size category. Segments represent the standard error of the means. ns indicates not significant.

Control treatments of the winter and spring genotypes showed different seed distribution across seed size categories (Fig. 2), where the winter genotype had larger seeds than the spring. Most of the seed size of winter control were recorded in the medium and large categories reaching 72.0 and 26.4%, respectively; while the spring control seeds mainly belonged to the small and medium categories, representing the 20.6 and 66.4% of total seeds, respectively (Fig. 2A). The S-S ratio reduction moved the seed size distribution to higher categories than the control in both winter and spring types of rapeseeds (Fig. 2). In the winter type, seed yield reductions of 63 and 79% in the small and medium categories, respectively, were recorded relative to the control (Table 1). On the contrary, this treatment increases 82% the large category and 392-fold the extreme-large category over the control. In the spring-type genotype, the S-S ratio treatment decreased the small category by 82% and 31% in the medium category, while the large category was increased 80-fold over the control (Table 1).

Seed categories of control treatments showed a peak at the 1.7-2.0 range for both genotypes (Fig. 2). The winter genotype had lower percentage in the 1.4-1.7 seed size category and higher percentage in the 2.0-2.36 range compared to the spring genotype (Fig. 2). The reduction of the S-S ratio led to the increase of seed size in both genotypes. The winter type showed higher seed distribution in the 2.0-2.36 range under lower S-S ratio, while the spring type maintained the peak in the 1.7-2.0 range (Fig. 2, Table 1). However, the seed proportion of the category 2.0-2.36 was increased in the spring type under the decreased S-S rate treatment.

When the seed number of the S-S ratio treatments was analysed at each category relative to the control in the winter type, a significant reduction of 80% was observed in the medium category, alongside a 9-fold increase in the very large category (Table 1). Additionally, the thousand-seed weight (TSW) showed an 11% reduction in the small category, and increases of 5%, 16%, and 20% in the medium, large, and very large categories, respectively (Table 1). In the spring type, there was an 82% reduction of seed number in the small category and a remarkable 67-fold increase in the large category. The TSW increased by 24% in the medium and 30% in the large categories (Table 1).

### 3.3 Seed oil, protein and macronutrients concentrations of the winter and spring-type genotypes under control and S-S ratio reduction

In the control winter-type genotype, seed size affected (p < 0.05) oil, protein, K, Ca, and Mg concentrations, while shading had impact (p < 0.05) on oil, protein (N) and S concentrations (Table 2). Interaction between seed size and S-S ratio was not found in this genotype. On the other hand, differences with the winter type were found in the spring genotype as the seed size in the control treatment showed effect on oil, P, K, S, Ca, and Mg concentrations, whereas the S-S ratio reduction impacted on oil and protein (N) concentrations (Table 2). In this genotype, interaction between S-S ratio and seed size was found for oil, protein, K, and S concentrations. Between winter and spring-type genotypes, differences were observed on protein (N), P, K, Ca, and Mg, with interactions between seed size and S-S ratio specifically for protein (N), K, and S (Table 2).

**Table 2.** Analysis of variance of seed oil and element concentrations of winter-type and spring-type rapeseed in response seed size categories and reduced source-sink ratio (S-S ratio) treatments.

| Source of variation | Oil | Macronutrients |  |  |  |  |  | Micronutrients |  |  |  |  | Na |
| --- | --- | --- | --- | --- | --- | --- | --- | --- | --- | --- | --- | --- | --- |
|  |  | N | P | K | S | Ca | Mg | B | Cu | Fe | Mn | Zn |  |
| Winter-type |  |  |  |  |  |  |  |  |  |  |  |  |  |
| Seed size category | ≤0.001 | 0.029 | 0.148 | ≤0.001 | 0.279 | ≤0.001 | 0.017 | 0.042 | 0.021 | 0.002 | 0.036 | 0.617 | ≤0.001 |
| S-S ratio | 0.002 | 0.009 | 0.858 | 0.211 | ≤0.001 | 0.279 | 0.540 | 0.042 | 0.009 | 0.113 | 0.377 | ≤0.001 | 0.006 |
| S x C | 0.474 | 0.454 | 0.687 | 0.183 | 0.182 | 0.679 | 0.258 | 0.228 | 0.827 | 0.839 | 0.023 | 0.584 | 0.059 |
| Spring-type |  |  |  |  |  |  |  |  |  |  |  |  |  |
| Seed size category | ≤0.001 | 0.124 | 0.009 | ≤0.001 | 0.032 | 0.003 | 0.004 | 0.003 | ≤0.001 | 0.853 | 0.701 | ≤0.001 | ≤0.001 |
| S-S ratio | 0.003 | 0.046 | 0.629 | 0.197 | 0.057 | 0.550 | 0.301 | ≤0.001 | ≤0.001 | 0.609 | 0.035 | ≤0.001 | 0.093 |
| S x C | 0.022 | ≤0.001 | 0.187 | ≤0.001 | 0.006 | 0.899 | 0.423 | 0.584 | 0.296 | 0.163 | 0.631 | ≤0.001 | 0.700 |
| Global |  |  |  |  |  |  |  |  |  |  |  |  |  |
| Type (T) | 0.379 | ≤0.001 | ≤0.001 | ≤0.001 | 0.826 | ≤0.001 | ≤0.001 | ≤0.001 | 0.012 | 0.378 | 0.076 | 0.102 | 0.351 |
| Size category (C) | ≤0.001 | 0.003 | 0.004 | ≤0.001 | 0.114 | ≤0.001 | ≤0.001 | ≤0.001 | 0.056 | 0.036 | 0.077 | 0.025 | ≤0.001 |
| S-S ratio (Type) (S(T)) | ≤0.001 | 0.002 | 0.899 | 0.289 | ≤0.001 | 0.403 | 0.518 | ≤0.001 | 0.001 | 0.327 | 0.072 | ≤0.001 | 0.002 |
| C x S(T) | 0.073 | 0.001 | 0.466 | 0.007 | 0.005 | 0.882 | 0.264 | 0.259 | 0.865 | 0.603 | 0.049 | 0.043 | 0.075 |

Significant p-values ( $p < 0.05$ ) are presented in bold type.

When oil, protein (N) and element concentrations were evaluated in the control treatments of the winter-type genotype, similar concentrations were found for oil, protein (N), P, and S across the seed size categories (Table 3). On the other hand, K, Ca, and Mg exhibited notable variations in small seeds (1.4-1.7 mm) compared to the other categories as K and Ca showed 19% and 28% higher seed concentrations than medium and large ones, respectively. On the contrary, Mg decreased by 8% (Table 3). Under the shading treatment, the small seed category reached 5% lower oil concentration than the other categories but higher concentration of K (24%), S (7%) and Ca (15%) within the large category (Table 3). When the S-S ratio effect was evaluated on oil and macronutrients concentrations relative to the control (Fig. S.1), the large size category showed lower oil (2%) and higher protein (9%) concentrations. The S-S ratio also increased seed concentration of S in the small (33%), medium (25%) and large (19%) seed categories. Moreover, Mg concentration decreased by only 6% in the medium category.

**Table 3.**
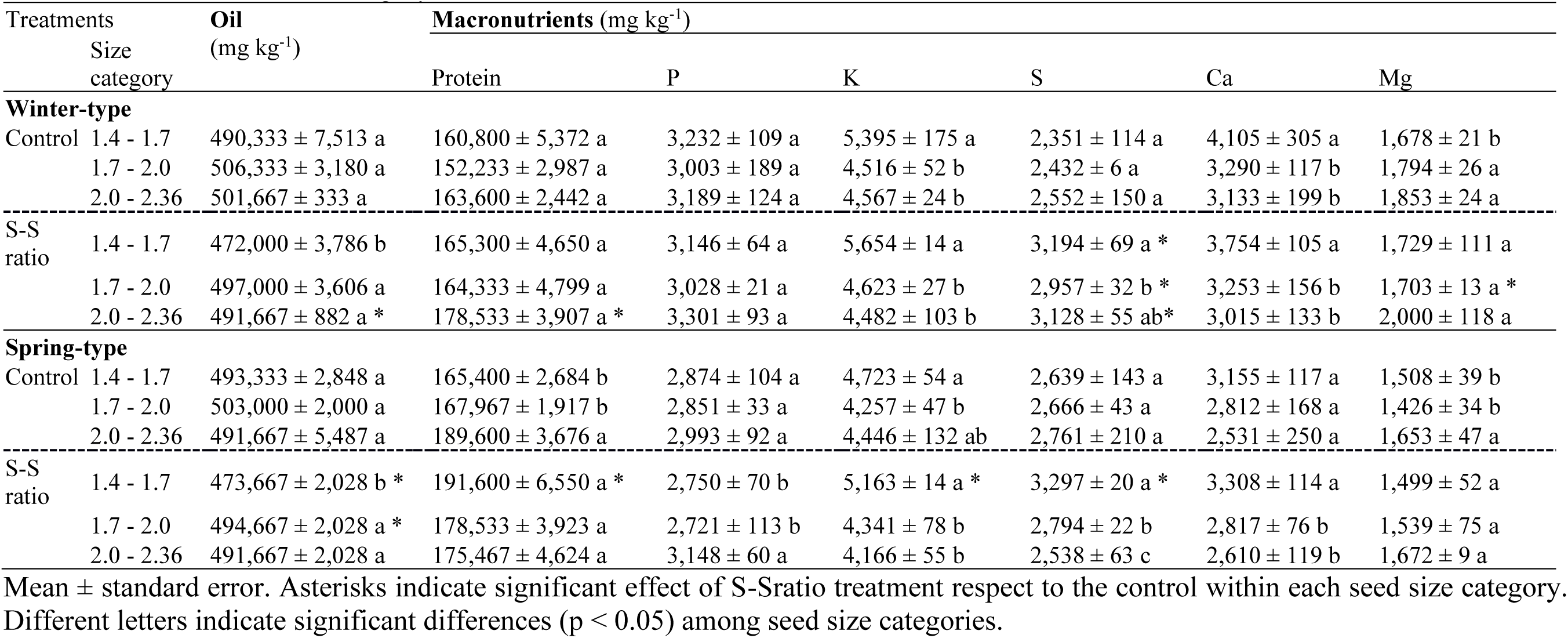
Seed oil and macronutrients concentrations of winter and spring rapeseed in control and reduced source-sink ratio (S-S ratio) treatments in each seed size category.

| Treatments |  | Oil<br>(mg kg<sup>-1</sup>) | Macronutrients (mg kg<sup>-1</sup>) |  |  |  |  |  |
| --- | --- | --- | --- | --- | --- | --- | --- | --- |
|  | Size<br>category |  | Protein | P | K | S | Ca | Mg |
| Winter-type |  |  |  |  |  |  |  |  |
| Control | 1.4 - 1.7 | 490,333 ± 7,513 a | 160,800 ± 5,372 a | 3,232 ± 109 a | 5,395 ± 175 a | 2,351 ± 114 a | 4,105 ± 305 a | 1,678 ± 21 b |
|  | 1.7 - 2.0 | 506,333 ± 3,180 a | 152,233 ± 2,987 a | 3,003 ± 189 a | 4,516 ± 52 b | 2,432 ± 6 a | 3,290 ± 117 b | 1,794 ± 26 a |
|  | 2.0 - 2.36 | 501,667 ± 333 a | 163,600 ± 2,442 a | 3,189 ± 124 a | 4,567 ± 24 b | 2,552 ± 150 a | 3,133 ± 199 b | 1,853 ± 24 a |
| S-S<br>ratio | 1.4 - 1.7 | 472,000 ± 3,786 b | 165,300 ± 4,650 a | 3,146 ± 64 a | 5,654 ± 14 a | 3,194 ± 69 a * | 3,754 ± 105 a | 1,729 ± 111 a |
|  | 1.7 - 2.0 | 497,000 ± 3,606 a | 164,333 ± 4,799 a | 3,028 ± 21 a | 4,623 ± 27 b | 2,957 ± 32 b * | 3,253 ± 156 b | 1,703 ± 13 a * |
|  | 2.0 - 2.36 | 491,667 ± 882 a * | 178,533 ± 3,907 a * | 3,301 ± 93 a | 4,482 ± 103 b | 3,128 ± 55 ab* | 3,015 ± 133 b | 2,000 ± 118 a |
| Spring-type |  |  |  |  |  |  |  |  |
| Control | 1.4 - 1.7 | 493,333 ± 2,848 a | 165,400 ± 2,684 b | 2,874 ± 104 a | 4,723 ± 54 a | 2,639 ± 143 a | 3,155 ± 117 a | 1,508 ± 39 b |
|  | 1.7 - 2.0 | 503,000 ± 2,000 a | 167,967 ± 1,917 b | 2,851 ± 33 a | 4,257 ± 47 b | 2,666 ± 43 a | 2,812 ± 168 a | 1,426 ± 34 b |
|  | 2.0 - 2.36 | 491,667 ± 5,487 a | 189,600 ± 3,676 a | 2,993 ± 92 a | 4,446 ± 132 ab | 2,761 ± 210 a | 2,531 ± 250 a | 1,653 ± 47 a |
| S-S<br>ratio | 1.4 - 1.7 | 473,667 ± 2,028 b * | 191,600 ± 6,550 a * | 2,750 ± 70 b | 5,163 ± 14 a * | 3,297 ± 20 a * | 3,308 ± 114 a | 1,499 ± 52 a |
|  | 1.7 - 2.0 | 494,667 ± 2,028 a * | 178,533 ± 3,923 a | 2,721 ± 113 b | 4,341 ± 78 b | 2,794 ± 22 b | 2,817 ± 76 b | 1,539 ± 75 a |
|  | 2.0 - 2.36 | 491,667 ± 2,028 a | 175,467 ± 4,624 a | 3,148 ± 60 a | 4,166 ± 55 b | 2,538 ± 63 c | 2,610 ± 119 b | 1,672 ± 9 a |

Mean ± standard error. Asterisks indicate significant effect of S-Sratio treatment respect to the control within each seed size category. Different letters indicate significant differences (p < 0.05) among seed size categories.

In the spring-type genotype, the control treatment did not affect the concentrations of oil, P, S, and Ca, but, impacted on protein (N), K, and Mg. Large seeds contained 14% more protein and 17% more Mg, while small seeds had a 9% higher K concentration relative to other categories (Table 4). Under shading conditions, the S-S treatment affected seed concentrations of oil, P, K, S, and Ca. In small seeds, oil concentration decreased by 4%, while K and Ca increased by 22% relative to other categories (medium and large). Conversely, large seeds showed a 13% rise in P concentration. Sulphur concentration varied across seed sizes, with small seeds exhibiting an 18% higher concentration, and large seeds 11% lower S than the medium size category (Table 4). When the decreased S-S ratio was assessed against the control treatment (Fig. S.1), oil concentration was lower in small (4%) and medium (2%) size seeds. However, small seeds showed higher protein (16%), K (11%), and S (27%) concentrations than in the control treatment seeds (Table 4).

**Table 4.**
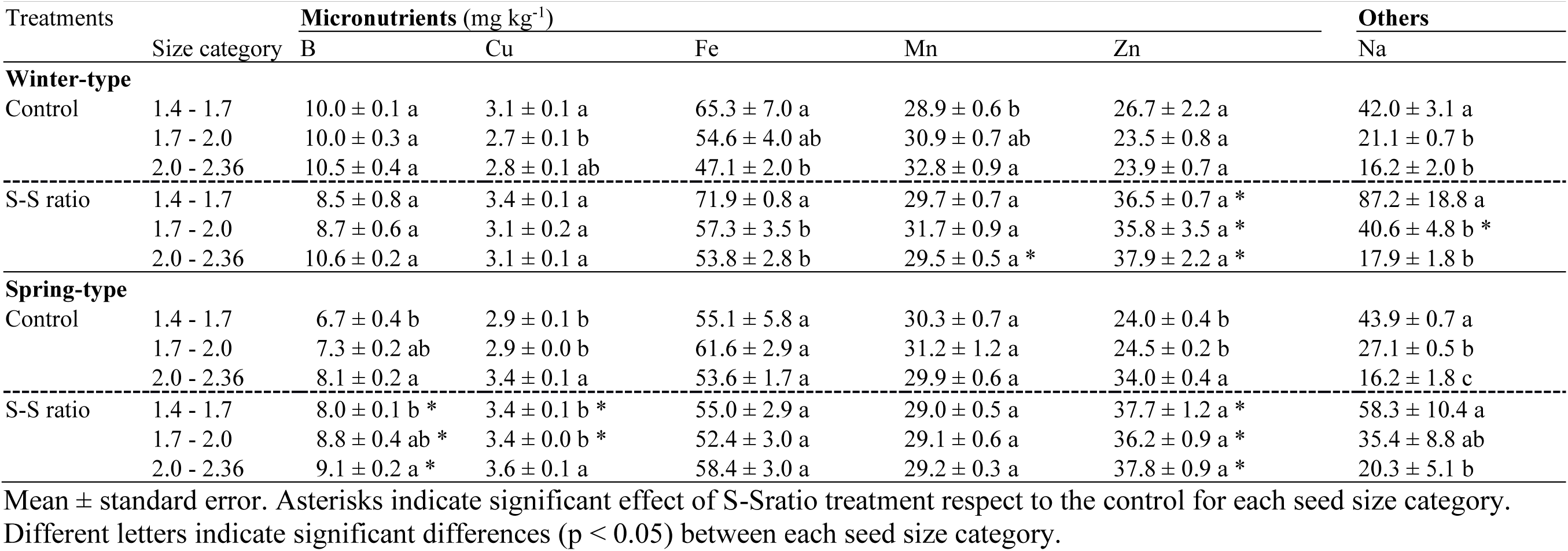
Seed micronutrients and sodium concentrations of winter and spring rapeseed in control and reduced source-sink ratio (S-S ratio) treatments in each seed size category.

| Treatments |  | Micronutrients (mg kg<sup>-1</sup>) |  |  |  |  | Others |
| --- | --- | --- | --- | --- | --- | --- | --- |
|  | Size category | B | Cu | Fe | Mn | Zn | Na |
| **Winter-type** |  |  |  |  |  |  |  |
| Control | 1.4 - 1.7 | 10.0 ± 0.1 a | 3.1 ± 0.1 a | 65.3 ± 7.0 a | 28.9 ± 0.6 b | 26.7 ± 2.2 a | 42.0 ± 3.1 a |
|  | 1.7 - 2.0 | 10.0 ± 0.3 a | 2.7 ± 0.1 b | 54.6 ± 4.0 ab | 30.9 ± 0.7 ab | 23.5 ± 0.8 a | 21.1 ± 0.7 b |
|  | 2.0 - 2.36 | 10.5 ± 0.4 a | 2.8 ± 0.1 ab | 47.1 ± 2.0 b | 32.8 ± 0.9 a | 23.9 ± 0.7 a | 16.2 ± 2.0 b |
| S-S ratio | 1.4 - 1.7 | 8.5 ± 0.8 a | 3.4 ± 0.1 a | 71.9 ± 0.8 a | 29.7 ± 0.7 a | 36.5 ± 0.7 a * | 87.2 ± 18.8 a |
|  | 1.7 - 2.0 | 8.7 ± 0.6 a | 3.1 ± 0.2 a | 57.3 ± 3.5 b | 31.7 ± 0.9 a | 35.8 ± 3.5 a * | 40.6 ± 4.8 b * |
|  | 2.0 - 2.36 | 10.6 ± 0.2 a | 3.1 ± 0.1 a | 53.8 ± 2.8 b | 29.5 ± 0.5 a * | 37.9 ± 2.2 a * | 17.9 ± 1.8 b |
| **Spring-type** |  |  |  |  |  |  |  |
| Control | 1.4 - 1.7 | 6.7 ± 0.4 b | 2.9 ± 0.1 b | 55.1 ± 5.8 a | 30.3 ± 0.7 a | 24.0 ± 0.4 b | 43.9 ± 0.7 a |
|  | 1.7 - 2.0 | 7.3 ± 0.2 ab | 2.9 ± 0.0 b | 61.6 ± 2.9 a | 31.2 ± 1.2 a | 24.5 ± 0.2 b | 27.1 ± 0.5 b |
|  | 2.0 - 2.36 | 8.1 ± 0.2 a | 3.4 ± 0.1 a | 53.6 ± 1.7 a | 29.9 ± 0.6 a | 34.0 ± 0.4 a | 16.2 ± 1.8 c |
| S-S ratio | 1.4 - 1.7 | 8.0 ± 0.1 b * | 3.4 ± 0.1 b * | 55.0 ± 2.9 a | 29.0 ± 0.5 a | 37.7 ± 1.2 a * | 58.3 ± 10.4 a |
|  | 1.7 - 2.0 | 8.8 ± 0.4 ab * | 3.4 ± 0.0 b * | 52.4 ± 3.0 a | 29.1 ± 0.6 a | 36.2 ± 0.9 a * | 35.4 ± 8.8 ab |
|  | 2.0 - 2.36 | 9.1 ± 0.2 a * | 3.6 ± 0.1 a | 58.4 ± 3.0 a | 29.2 ± 0.3 a | 37.8 ± 0.9 a * | 20.3 ± 5.1 b |

Mean ± standard error. Asterisks indicate significant effect of S-Sratio treatment respect to the control for each seed size category.
Different letters indicate significant differences (p < 0.05) between each seed size category.

### 3.4 Micronutrients and sodium concentrations of winter-type and spring-type rapeseed

The seed size modified (p < 0.05) seed concentration of B, Cu, Fe, Mn, and Na of the winter-type genotype (Table 2). In addition, the shading treatment affected B, Cu, Zn, and Na seed concentrations. Interaction between seed size and S-S ratio treatments was only found in Mn. On the other hand, in the spring-type genotype, seed size affected B, Cu, Zn, and Na, while the S-S ratio impacted on B, Cu, Mn, and Zn seed concentrations (Table 2). The interaction between seed size and S-S ratio treatments was only recorded for Zn. Comparing winter and spring-types, differences were observed for B and Cu, with interactions between seed size and S-S ratio in Mn and Zn.

Seed size of control winter-type genotype reported similar concentrations of B and Zn, while differences were noted for Cu, Fe, Mn, and Na (Table 4). Specifically, Cu was 15% higher in large seeds compared to small ones, while Fe was 39% higher and Mn 6% lower in small seeds relative to large seeds. Sodium concentrations were almost double (99%) in small seeds compared to the other categories. When the effect of the decreased S-S ratio was analysed, the size of the seeds showed similar values of B, Cu, Mn, and Zn concentrations. Small seeds exhibited 29% higher Fe and 119% higher Na concentrations relative to other categories, i.e. medium and large size (Table 4).

In the control spring-type genotype, seed size had specific effects on micronutrients B, Zn, and Na. Boron concentration was 21% higher in large seeds compared to small ones, while Cu and Zn were 17 and 40% higher, respectively, in large seeds than in the other seed size categories. Sodium was 62% higher in small seeds and 40% lower in large than in medium seeds (Table 4). Under shading, seed size did not modify seed concentrations of Fe, Mn, and Zn, but this S-S ratio treatment affected B, Cu, and Na concentrations (Fig. S.1). In large seeds, B over-yielded small seeds by 14%, and also in Na (64%) and Cu (6%) (Table 5).

When the shading treatment was considered, the decreased S-S ratio improved B, Zn, and Cu concentrations by 19, 21 and 12% of the small, medium, and large seeds, respectively, over the control seeds (Figure S.1). Zinc concentrations rose by 57%, 48%, and 11%, while copper increased by 17% in both the small and medium seed categories.

### 3.5 Relationships among seed size, weight and quality traits

An integrated PCA analysis was conducted to assess associations among yield, its numerical components, and quality traits in response to the seed size and S-S treatments of winter and spring rapeseed genotypes (Fig. 3). Components PC1 and PC2 accounted for 60% of the total variation, capturing the dataset variability. PC1 primarily differentiated yield, yield components and oil concentration from most of seed elements concentrations, while PC2 mostly captured variations in element concentrations, specifically Ca, K, and Na, in contrast to protein (N) and other elements such as Cu and Zn. However, there was not clear separation between different seed size categories and S-S ratio treatments (Fig. 3).

**Figure 3.**
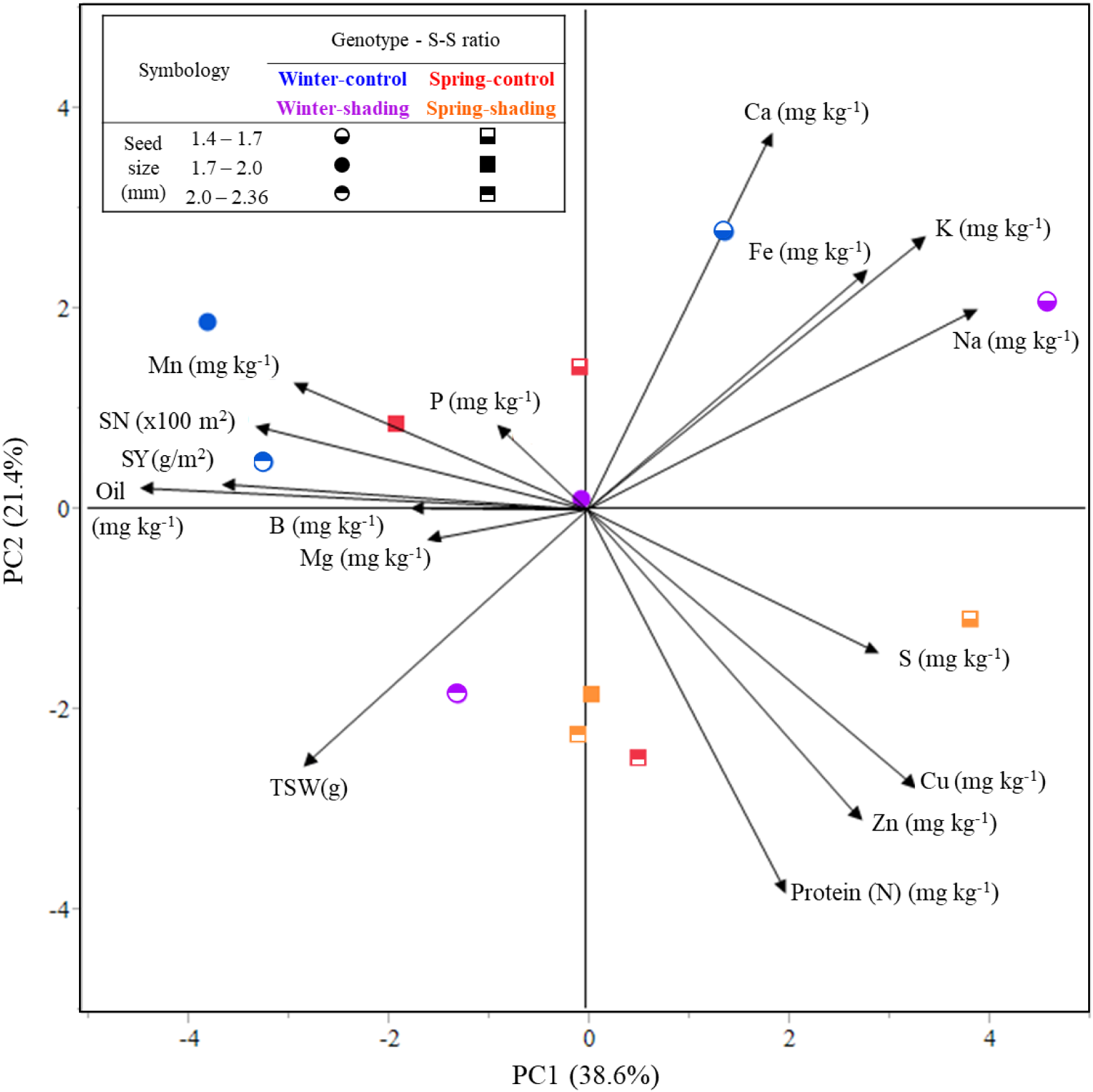
Principal component analysis (PCA) of seed yield (SY), seed number (SN), thousand seed weight (TSW), oil and nutrients concentrations of winter-type (circles symbols) and spring-type (squares symbols) genotypes in control (blue and red symbols) and decreased S-S ratio (purple and orange symbols) treatments of seed size categories: small (lower half closed symbols), medium (closed symbols), and large (upper half closed symbols) categories.

Noteworthy, seed yield, seed number, seed oil and Mn concentrations were grouped together in the winter type. Showing no trade-off between oil concentration and seed yield and oil. Nevertheless, this group showed negative association with another group gathering protein (N), K, S, Cu and Zn (Fig. 3), supporting a trade-off between seed yield and oil concentration with key nutrients such as protein, S, Zn and Cu. TSW was positively correlated with Mg and B, but negatively associated with K, Ca, Fe, and Na (Fig. 4). Thus, larger seeds achieved higher concentrations of Mg and B, while had lower concentrations of K, Ca, Fe, and Na.

**Figure 4.**
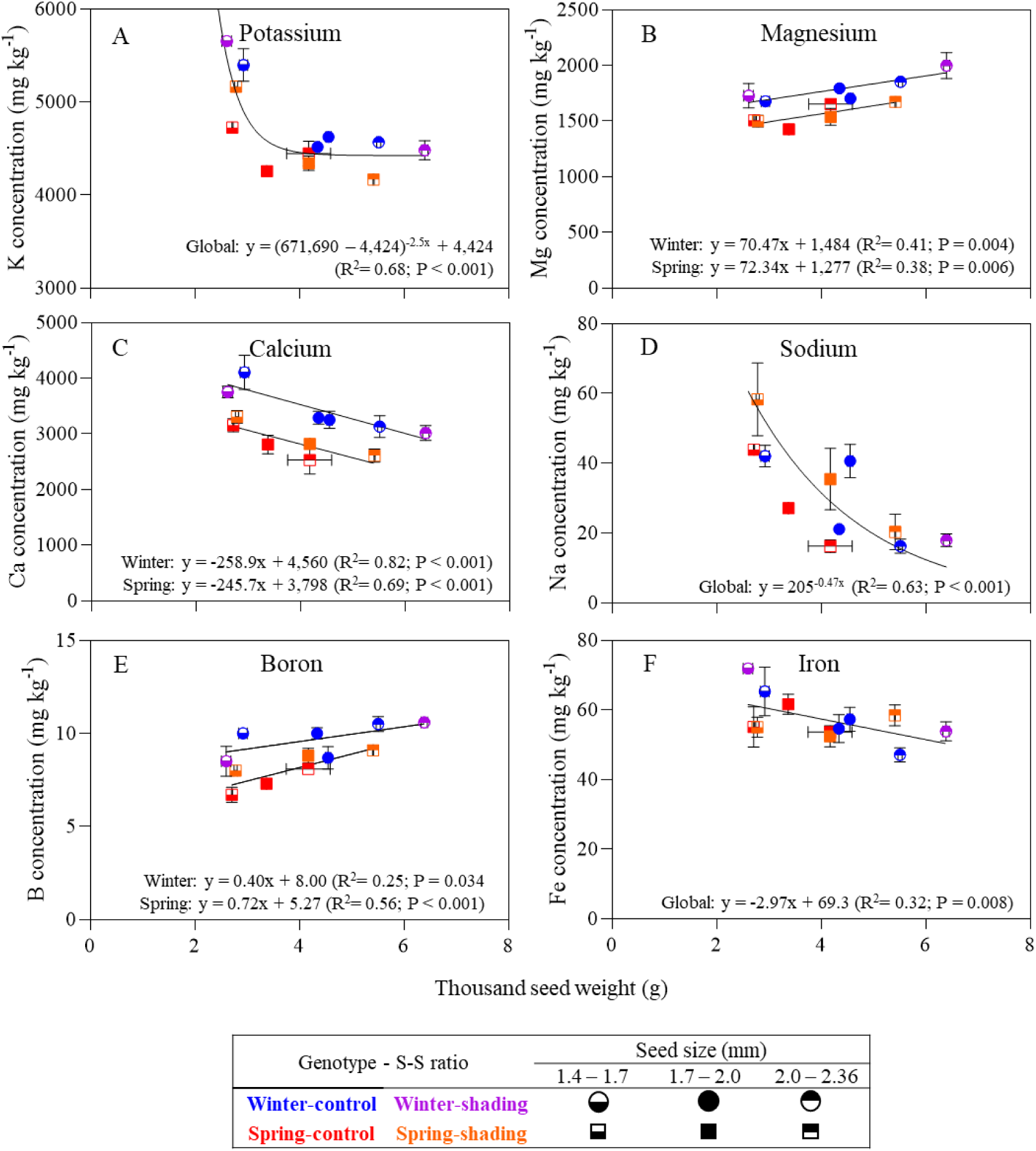
Relationships between seed concentrations of potassium (A), magnesium (B), calcium (C), sodium (D), boron (E) or iron (F) and thousand seed weight (TSW) of winter-type (circle symbols) and spring-type (square symbols) genotypes in control (blue and red symbols) and S-S ratio (purple and orange symbols) treatments ofs seed size categories: small (lower half closed symbols), medium (closed symbols), and large (upper half closed symbols) categories.

In addition to the PCA analysis, the associations between specific seed element concentrations and TSW were evaluated across the seed size categories (Fig. 4). K and Na exhibited a one-phase decay function, while Mg, Ca, and B followed a linear regression with different functions between winter and spring-type genotypes. The winter-type genotypes showed higher concentrations of these elements compared to the spring-type. It is worth noting that K concentration decreased as seed size increased, reaching a plateau of approximately 4,424 mg K kg^-1^ in seeds exceeding TSW by 3.37 g.

## 4. Discussion

### 4.1 Sensitivity of quality traits to seed size

This study analyses the effect of seed size categories, as well as the impact of the S-S reduction at the beginning of seed filling on key quality traits (seed concentrations of oil, protein, macronutrients and micronutrients) of winter and spring rapeseed genotypes. Seed material used in this study were collected from a previous work by Verdejo and Calderini (2020) from control plots and those showing significant difference of seed weight due to lower S-S ratio treatments. Oil, protein and elements concentrations in seeds of rapeseed are of central importance for this crop for food and feed industries (Fobert et al., 2008; Thiyam-Hollaender et al., 2013; Vahedifar and Wu, 2022). Previous studies focused on oil and protein seed concentrations of this crop in experiments were seed yield ranged between 2.43 and 5.5 Mg ha^-1^, reaching between 38.9% and 51.9% oil and from 14.3% to 37.2% protein concentrations under controlled and field conditions (Diepenbrock, 2000; Labra et al., 2017; Kirkegaard et al., 2018; Verdejo and Calderini, 2020). We aimed to extend the information to high-yield environments such as that of southern Chile, and similar environments reaching seed yields of rapeseed by 6.5 Mg ha^-1^ or more, together with ∼51% seed oil concentration (Berry and Spink, 2006; Verdejo and Calderini, 2020; Rivelli et al., 2024). Even more important, it is little known the effect of both seed size and the source-sink ratio on quality traits of winter and spring rapeseed genotypes as only partial information has been reported in this regard.

Remarkably, seed quality trait concentrations measured here are not independent of seed yield, as previous studies showed interactions between nutrient concentration and crop yield (McDonald et al., 2008; Rondanini et al., 2014; Richard et al., 2025). Although earlier works focused on seed yield and oil and protein concentration, often under controlled conditions and specific perturbations (treatments), our study extends those traits to include macro- and micronutrients concentrations in seeds. Seed yield and seed weight have shown resilience to environmental stresses (Labra et al., 2017; Kirkegaard et al., 2018; Verdejo and Calderini, 2020; Rivelli et al., 2024), however, it remains unclear whether this resilience also is true for seed elements concentrations and the sensitivity of seed oil, protein and element concentrations in different seed sizes of winter and spring rapeseed genotypes.

### 4.2 Seed size on oil and element concentrations of winter-type and spring-type rapeseed

The selected seed size categories were chosen accounting for comparisons with a previous study evaluating a single spring rapeseed genotype (Beyzi et al., 2019). In our experiments, both total TSW and percentages of seed size categories showed differences between winter and spring genotypes, where the winter type reached higher values than the spring one in control and S-S ratio treatments. In control plots, seeds belonging to the medium category (mean ± standard deviation) accounted for 70.8% of seeds across winter and spring-type genotypes. The high seed proportion between 1.7 and 2.0 mm has often been reported in the literature (Çalışır et al., 2005; Beyzi et al., 2019; Li et al., 2019). Oil seed concentrations of winter and spring-type genotypes 0in this study exhibited higher concentration as reported by Beyzi et al. (2019), but no information was reported by the authors on protein (N) seed concentration. Between oil and protein concentrations, a trade-off has been commonly described in the literature (Rondanini et al., 2014; Labra et al., 2017; Kirkegaard et al., 2018). Comparing studies from previous research conducted in southern Chile, oil concentration is higher and more stable than protein, in rapeseed and even in sunflower (Castillo et al., 2017; Labra et al., 2017; Rivelli et al., 2024). For example, the winter-type exhibited 2% lower oil and 9% higher protein concentrations in the large seed size category, while in the spring-type, lower oil seed concentrations were recorded in small (4%) and medium (2%) seed sizes, accompanied by a 16% increase in protein concentration in small seeds.

The others key quality traits for food and feed of rapeseed are the elements seed concentrations. Beyzi et al. (2019) reported, higher concentrations of P, K, Ca, Mg, Cu, Fe, Mn, Zn, and Na, along with lower concentrations of S and B than the present study across different seed sizes. Higher seed elemental concentrations compared to our findings were found also in other rapeseed studies (Çalışır et al., 2005; Musa Özcan, 2006). These differences may be attributed to the origin of the plant material, as Beyzi et al. (2019) worked with seeds planned for sowing, while our study focused on seeds for oil extraction. This consideration is important because the seed type and its intended use influence the results and applicability of the findings. No less important is seed yield reached across the different studies, as a dilutions effect on element concentration is often found under high seed yield (McDonald et al., 2008). Seed yield reached in our study was 8.4 Mg ha^-1^ in winter and 6.6 Mg ha^-1^ in spring type genotypes. These can significantly impact element concentrations. Additionally, the high oil seed concentration values reported in this study may influence the element seed, reducing its concentrations (Newkirk, 2011) .

In the present study, element concentrations responded differently across genotypes. In control plots, winter-type rapeseed showed variation in K, Ca, and Mg across seed sizes, whereas the spring-type exhibited variation in protein, K, and Mg. Following the same pattern, K concentration was higher, and Mg concentration was lower in small seeds of both genotypes; however, no correlation was observed between the two nutrients. This pattern contrasts with Beyzi et al. (2019), since they found lower concentrations of K and Mg in small seeds compared to larger seeds. Regarding Na, our results showed relatively low concentrations (ranging from 16.2 to 87.2 mg kg^-1^) compared to values reported in other studies, which can reach up to 202 mg kg^-1^ (Beyzi et al., 2019). This low Na content is advantageous for both human and animal nutrition, as excessive sodium intake is linked to hypertension and cardiovascular risks in humans (Grillo et al., 2019), and can disrupt the electrolyte balance required for optimal physiological performance in livestock (Goff, 2018)(Doyle et al., 2022). In our study, S concentrations of the winter-type increase by 33%, 25%, and 19% in the small, medium, and large categories, respectively, while the spring-type showed a 27% increase in small seeds. The rise in S concentration is of significant interest due to the importance of S for food, feed and seed germination (Massuia de Almeida et al., 2021; Mondal et al., 2022; Rasheed et al., 2024). The correlations between quality traits and both yield and yield components underscore the complex interactions among genotype, seed size, and environmental conditions, highlighting their significance for rapeseed cultivation practices.

### 4.3 Source-sink reduction on seed element concentrations of winter-type and spring-type rapeseed

The evaluation of the S-S ratio reduction on seed size and quality traits is supported by the fact that rapeseed is a very plastic species, where changes in seed yield components could compensate for one another. For example, when SN was affected by the S-S ratio reduction, TSW totally or partially compensated SN decrease (Labra et al., 2017; Kirkegaard et al., 2018; Verdejo and Calderini, 2020; Rivelli et al., 2024). Changes in SN also occur in farmers’ conditions, mainly if a biotic or abiotic constrain affect SN setting during the critical period of rapeseed (from 100 to 500°Cd after flowering), with the concomitant effect of increasing TSW (Kirkegaard et al., 2018). The effect of the S-S ratio modified the seed size proportions in both winter and spring types of genotypes, highlighting the importance of evaluating whether this modification alters the oil and element concentrations of rapeseed. However, other abiotic stresses, such as heat stress treatments during grain filling, have little effect on the seed weight of rapeseed (Verdejo and Calderini, 2025). Currently, Rivelli et al. (2021) has reported that an increase in daytime temperature and a reduction of incident solar radiation already happened in the Southern Cone of South America throughout the last 50 years. Therefore, there is an urgent need to know the effects of reductions in the source-sink ratio on rapeseed yield and seed quality traits. Since shading treatments affected the seed distribution across sizes in both types by increasing the proportion of large seeds, it is important to note that, although differences in oil and element concentrations were observed, most of the element contents (g m^-2^) showed an overall decrease due to reduced S-S ratio treatment (Tables A.1 and A.2).

## 5. Conclusions

Seed oil and protein concentrations exhibited differential responses based on genotype, seed size, and S-S ratio treatment. Smaller seeds generally demonstrated reduced oil concentration alongside the increased protein content. Specifically, under shading conditions, the small seed category showed a 5% lower oil concentration, while seed protein concentration increased by 6% in both genotypes. In the winter type, large seeds under the S-S ratio reduction, presented 2% less oil and 9% higher protein concentration. Similarly, in the spring genotype, small and medium seeds had 4% and 2% lower oil concentrations, respectively, while small seeds showed a 16% increase in protein. Notably, oil concentration was positively correlated with seed yield and negatively associated with protein. Concerning elements’ composition, distinct effects were observed depending on seed size and genotype. Large seeds generally exhibited higher Mg and B concentrations, but lower levels of K, Ca, Fe and Na. In the control winter genotype, small seeds showed 19% and 28% higher K and Ca concentrations, respectively, but 8% lower Mg compared to other size categories. Under shading, winter small seeds recorded higher concentrations of K (24%), S (7%), and Ca (15%). For the spring genotype, S-S ratio reduction led to a 22% increase in K and Ca, and a 27% increase in S in small seeds, while large seeds showed a 13% rise in P. Furthermore, oil concentration was negatively associated with the concentrations of K, S, Cu, and Zn. These findings highlight the complex interactions between seed size, genotype, and environmental conditions, and their collective impact on the nutrient profiles of rapeseed. This information is helpful for optimising cultivation practices and breeding strategies.

## 6. Author contribution

José F. Verdejo: Conceptualization, data curation, formal analysis, investigation, methodology, visualization, writing - original draft. Daniel F. Calderini: Conceptualization, investigation, methodology, resources, supervision, writing – review and editing. All authors read and approved the final version of the manuscript.

## Acknowledgements

We thank Olaff Sass (NPZ-Lembke, Germany) and Erik von Baer (Semillas Baer, Chile) for kindly providing the seeds of the genotypes. We really appreciate technical support from the staff of the Austral Farming Experimental Station (EEAA) of the Universidad Austral de Chile. The present study was supported by Project FONDECYT 1170913 (Chilean Technical and Scientific Research Council, CONICYT/ANID) competitive grant. José Verdejo held a postgraduate scholarship from CONICYT (ANID) 2017-21171384.

---

**Figure A.1.**
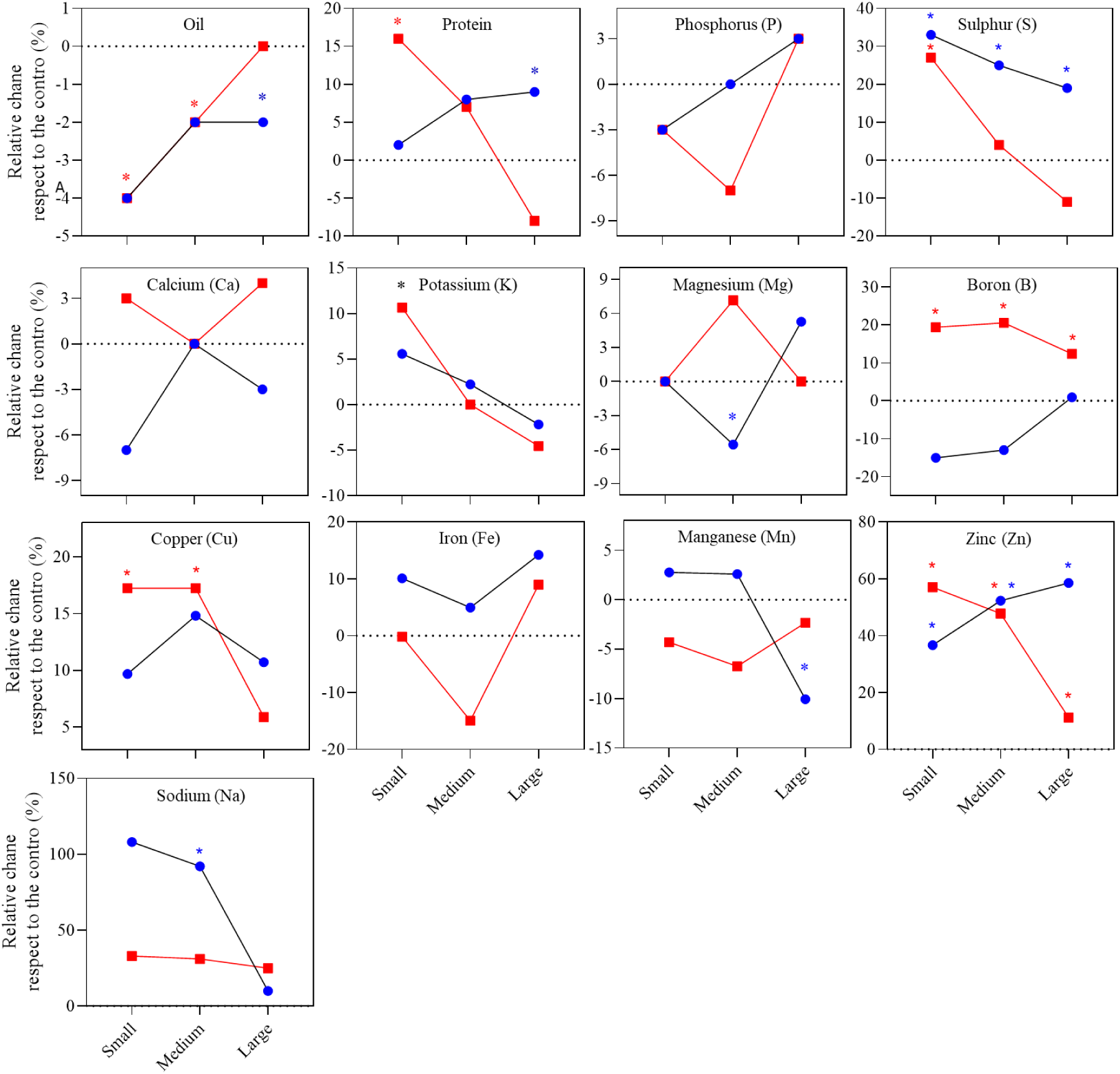
Relative change of the element concentration due to the S-S ratio respect to the control of winter (blue squares) and spring rapeseed (red circles). Asterisks indicate a significant effect of the S-S ratio treatment compared to the control for each seed size category.

**Table A.1.**
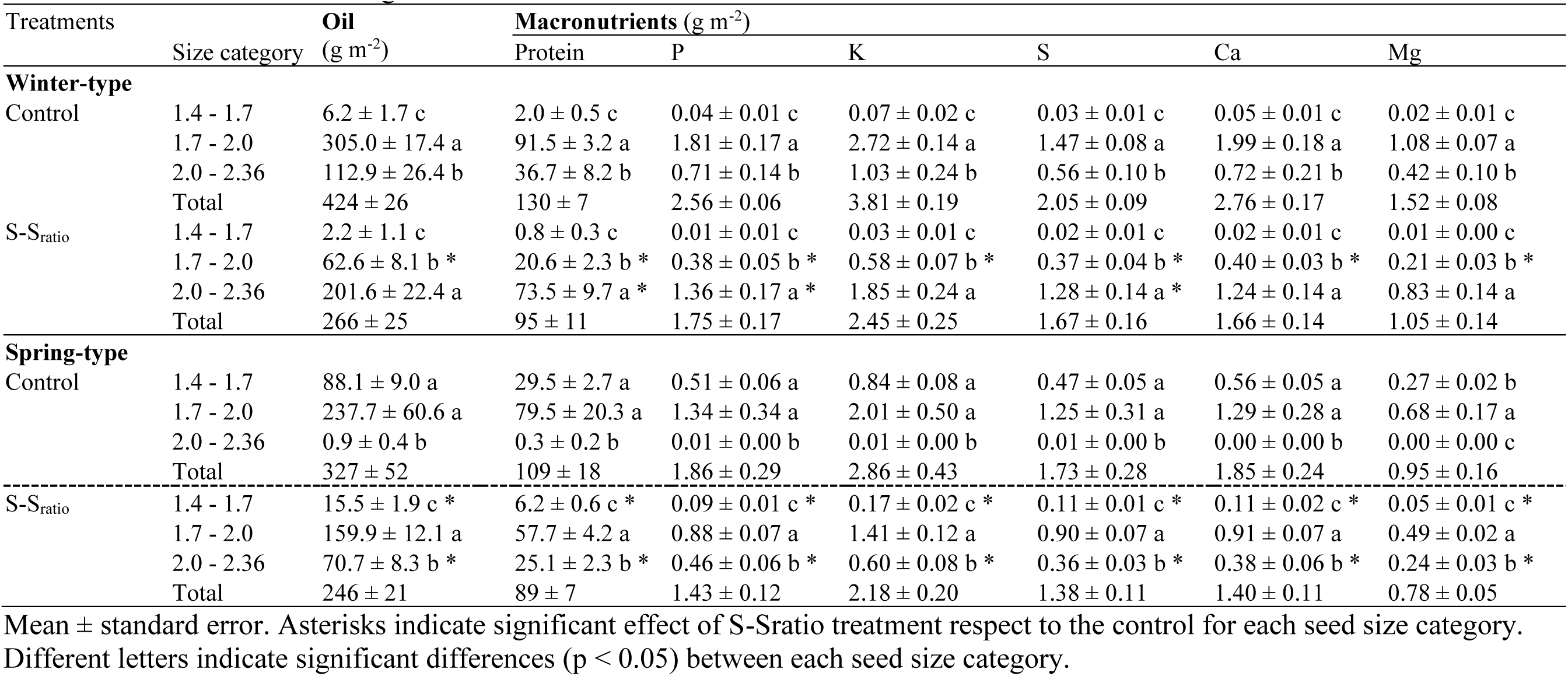
Seed oil and macronutrients content of winter and spring rapeseed in control and reduced source-sink ratio (S-S ratio) treatments in each seed size categories.

| Treatments in each seed size category: |  |  |  |  |  |  |  |  |
| --- | --- | --- | --- | --- | --- | --- | --- | --- |
| Treatments |  | Oil | Macronutrients (g m<sup>-2</sup>) |  |  |  |  |  |
|  | Size category | (g m<sup>-2</sup>) | Protein | P | K | S | Ca | Mg |
| **Winter-type** |  |  |  |  |  |  |  |  |
| Control | 1.4 - 1.7 | 6.2 ± 1.7 c | 2.0 ± 0.5 c | 0.04 ± 0.01 c | 0.07 ± 0.02 c | 0.03 ± 0.01 c | 0.05 ± 0.01 c | 0.02 ± 0.01 c |
|  | 1.7 - 2.0 | 305.0 ± 17.4 a | 91.5 ± 3.2 a | 1.81 ± 0.17 a | 2.72 ± 0.14 a | 1.47 ± 0.08 a | 1.99 ± 0.18 a | 1.08 ± 0.07 a |
|  | 2.0 - 2.36 | 112.9 ± 26.4 b | 36.7 ± 8.2 b | 0.71 ± 0.14 b | 1.03 ± 0.24 b | 0.56 ± 0.10 b | 0.72 ± 0.21 b | 0.42 ± 0.10 b |
|  | Total | 424 ± 26 | 130 ± 7 | 2.56 ± 0.06 | 3.81 ± 0.19 | 2.05 ± 0.09 | 2.76 ± 0.17 | 1.52 ± 0.08 |
| S-S<sub>ratio</sub> | 1.4 - 1.7 | 2.2 ± 1.1 c | 0.8 ± 0.3 c | 0.01 ± 0.01 c | 0.03 ± 0.01 c | 0.02 ± 0.01 c | 0.02 ± 0.01 c | 0.01 ± 0.00 c |
|  | 1.7 - 2.0 | 62.6 ± 8.1 b * | 20.6 ± 2.3 b * | 0.38 ± 0.05 b * | 0.58 ± 0.07 b * | 0.37 ± 0.04 b * | 0.40 ± 0.03 b * | 0.21 ± 0.03 b * |
|  | 2.0 - 2.36 | 201.6 ± 22.4 a | 73.5 ± 9.7 a * | 1.36 ± 0.17 a * | 1.85 ± 0.24 a | 1.28 ± 0.14 a * | 1.24 ± 0.14 a | 0.83 ± 0.14 a |
|  | Total | 266 ± 25 | 95 ± 11 | 1.75 ± 0.17 | 2.45 ± 0.25 | 1.67 ± 0.16 | 1.66 ± 0.14 | 1.05 ± 0.14 |
| **Spring-type** |  |  |  |  |  |  |  |  |
| Control | 1.4 - 1.7 | 88.1 ± 9.0 a | 29.5 ± 2.7 a | 0.51 ± 0.06 a | 0.84 ± 0.08 a | 0.47 ± 0.05 a | 0.56 ± 0.05 a | 0.27 ± 0.02 b |
|  | 1.7 - 2.0 | 237.7 ± 60.6 a | 79.5 ± 20.3 a | 1.34 ± 0.34 a | 2.01 ± 0.50 a | 1.25 ± 0.31 a | 1.29 ± 0.28 a | 0.68 ± 0.17 a |
|  | 2.0 - 2.36 | 0.9 ± 0.4 b | 0.3 ± 0.2 b | 0.01 ± 0.00 b | 0.01 ± 0.00 b | 0.01 ± 0.00 b | 0.00 ± 0.00 b | 0.00 ± 0.00 c |
|  | Total | 327 ± 52 | 109 ± 18 | 1.86 ± 0.29 | 2.86 ± 0.43 | 1.73 ± 0.28 | 1.85 ± 0.24 | 0.95 ± 0.16 |
| S-S<sub>ratio</sub> | 1.4 - 1.7 | 15.5 ± 1.9 c * | 6.2 ± 0.6 c * | 0.09 ± 0.01 c * | 0.17 ± 0.02 c * | 0.11 ± 0.01 c * | 0.11 ± 0.02 c * | 0.05 ± 0.01 c * |
|  | 1.7 - 2.0 | 159.9 ± 12.1 a | 57.7 ± 4.2 a | 0.88 ± 0.07 a | 1.41 ± 0.12 a | 0.90 ± 0.07 a | 0.91 ± 0.07 a | 0.49 ± 0.02 a |
|  | 2.0 - 2.36 | 70.7 ± 8.3 b * | 25.1 ± 2.3 b * | 0.46 ± 0.06 b * | 0.60 ± 0.08 b * | 0.36 ± 0.03 b * | 0.38 ± 0.06 b * | 0.24 ± 0.03 b * |
|  | Total | 246 ± 21 | 89 ± 7 | 1.43 ± 0.12 | 2.18 ± 0.20 | 1.38 ± 0.11 | 1.40 ± 0.11 | 0.78 ± 0.05 |

Mean ± standard error. Asterisks indicate significant effect of S-Sratio treatment respect to the control for each seed size category.
Different letters indicate significant differences (p < 0.05) between each seed size category.

**Table A.2.**
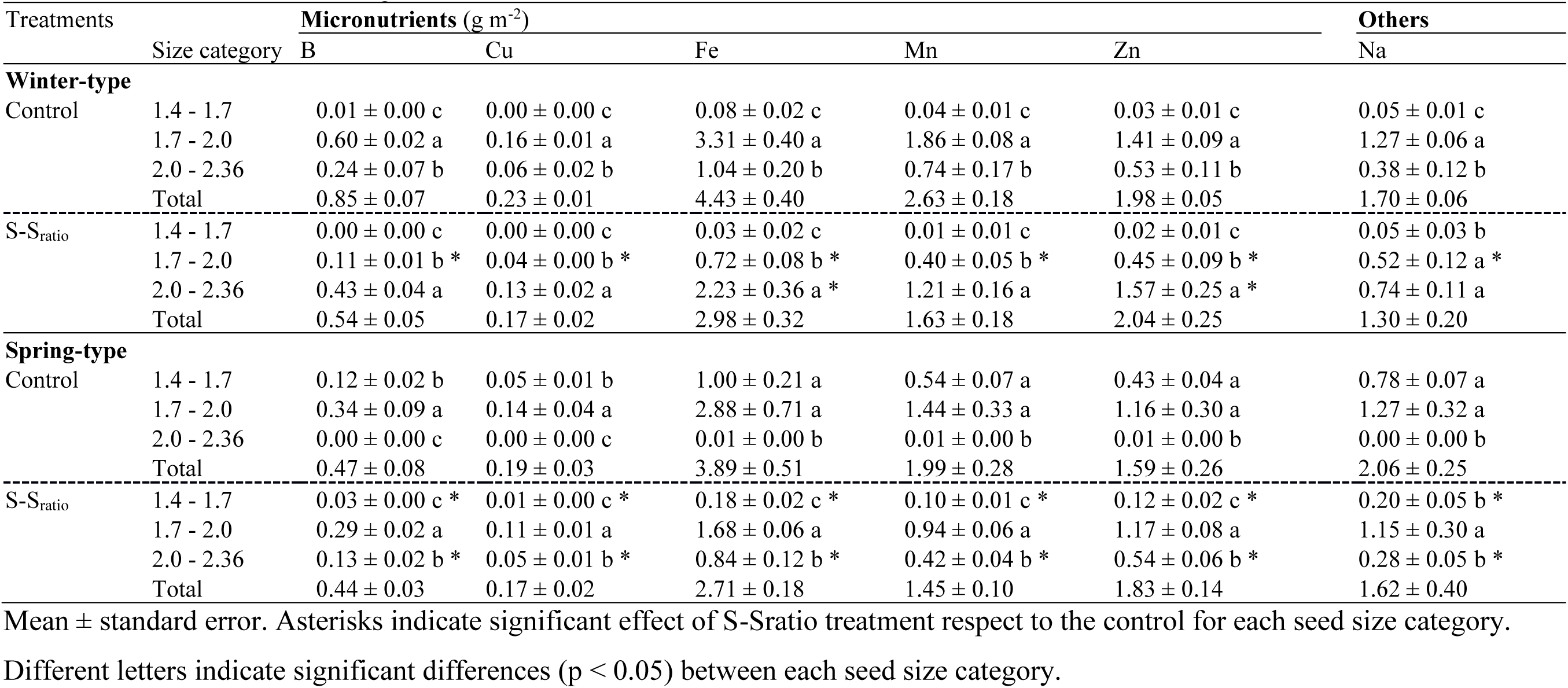
Seed micronutrients and sodium content of winter and spring rapeseed in control and reduced source-sink ratio (S-S ratio) treatments in each seed size categories.

## References

1. Batool, M., El-Badri, A.M., Wang, C., Mohamed, I.A.A., Wang, Z., Khatab, A., Bashir, F., Xu, Z., Wang, J., Kuai, J., Wang, B., Zhou, G., 2022. The role of storage reserves and their mobilization during seed germination under drought stress conditions of rapeseed cultivars with high and low oli contents. Crop and Environment 1, 231–240. 10.1016/j.crope.2022.09.003

2. Berry, P.M., Spink, J.H., 2006. A physiological analysis of oilseed rape yields: Past and future. The Journal of Agricultural Science 144, 381–392. 10.1017/S0021859606006423

3. Beyzi, E., Gunes, A., Buyukkilic Beyzi, S., Konca, Y., 2019. Changes in fatty acid and mineral composition of rapeseed (Brassica napus ssp. oleifera L.) oil with seed sizes. Ind Crop Prod 129, 10–14. 10.1016/j.indcrop.2018.11.064

4. Calderini, D.F., Ortiz-Monasterio, I., 2003. Grain Position Affects Grain Macronutrient and Micronutrient Concentrations in Wheat. Crop Sci 43, 141–151. 10.2135/cropsci2003.1410

5. Çalışır, S., Marakoğlu, T., Öğüt, H., Öztürk, Ö., 2005. Physical properties of rapeseed (Brassica napus oleifera L.). Journal of Food Engineering 69, 61–66. 10.1016/j.jfoodeng.2004.07.010

6. Castillo, F.M., Vásquez, S.C., Calderini, D.F., 2017. Does the pre-flowering period determine the potential grain weight of sunflower? Field Crop Res 212, 23–33. 10.1016/j.fcr.2017.06.029

7. del Pozo, A., Jobet, C., Matus, I., Méndez-Espinoza, A.M., Garriga, M., Castillo, D., Elazab, A., 2022. Genetic yield gains and changes in morphophysiological-related traits of winter wheat in southern chilean high-yielding environments. Front Plant Sci 12. 10.3389/fpls.2021.732988

8. Diepenbrock, W., 2000. Yield analysis of winter oilseed rape (Brassica napus L.): a review. Field Crop Res 67, 35–49. 10.1016/S0378-4290(00)00082-4

9. FAO, IFAD, UNICEF, WFP, WHO, 2023. The State of Food Security and Nutrition in the World 2023. Urbanization, agrifood systems transformation and healthy diets across the rural–urban continuum. FAO, Rome, Italy.

10. Farsak, H., 2009. The Effect of Different Row Spacing on the Yield and Yield Components of Rapeseed Varietes. In: Secondary Farsak, H. (Ed.), Secondary The Effect of Different Row Spacing on the Yield and Yield Components of Rapeseed Varietes. Publisher, Aydin p.^pp. Pages

11. Fobert, P.R., Smith, M.A., Zou, J., Mietkiewska, E., Keller, W.A., Taylor, D.C., 2008. Developing Canadian seed oils as industrial feedstocks. Biofuel Bioprod Bior 2, 206–214. 10.1002/bbb.72

12. Goff, J.P., 2018. Invited review: Mineral absorption mechanisms, mineral interactions that affect acid-base and antioxidant status, and diet considerations to improve mineral status. J Dairy Sci 101, 2763–2813. 10.3168/jds.2017-13112

13. Grillo, A., Salvi, L., Coruzzi, P., Salvi, P., Parati, G., 2019. Sodium Intake and Hypertension. Nutrients 11, 1970. 10.3390/nu11091970

14. Gulden, R.H., Thomas, A.G., Shirtliffe, S.J., 2004. Relative contribution of genotype, seed size and environment to secondary seed dormancy potential in Canadian spring oilseed rape (Brassica napus). Weed Research 44, 97–106. 10.1111/j.1365-3180.2003.00377.x

15. Hefferon, K.L., 2015. Nutritionally enhanced food crops; progress and perspectives. Int J Mol Sci 16, 3895–3914. 10.3390/ijms16023895

16. Jensen, S.K., Liu, Y.-G., Eggum, B.O., 1995. The influence of seed size and hull content on the composition and digestibility of rapeseeds in rats. Anim Feed Sci Tech 54, 9–19. 10.1016/0377-8401(94)00762-X

17. Kirk, P.L., 1950. Kjeldahl method for total nitrogen. Analytical Chemistry 22, 354–358. 10.1021/ac60038a038

18. Kirkegaard, J.A., Lilley, J.M., Brill, R.D., Ware, A.H., Walela, C.K., 2018. The critical period for yield and quality determination in canola (Brassica napus L.). Field Crop Res 222, 180–188. 10.1016/j.fcr.2018.03.018

19. Kowalska, G., Kowalski, R., Hawlena, J., Rowiński, R., 2020. Seeds of oilseed rape as an alternative source of protein and minerals. J Elem 25, 513–522. 10.5601/jelem.2019.24.3.1893

20. Kreitzman, M., Toensmeier, E., Chan, K.M.A., Smukler, S., Ramankutty, N., 2020. Perennial Staple Crops: Yields, Distribution, and Nutrition in the Global Food System. Frontiers in Sustainable Food Systems 4. 10.3389/fsufs.2020.588988

21. Kutner, M., Nachtsheim, C., Neter, J., 2004. Applied Linear Regression Models. McGraw-Hill Education, Boston.

22. Labra, M.H., Struik, P.C., Evers, J.B., Calderini, D.F., 2017. Plasticity of seed weight compensates reductions in seed number of oilseed rape in response to shading at flowering. Eur J Agron 84, 113–124. 10.1016/j.eja.2016.12.011

23. Lamb, K.E., Johnson, B.L., 2004. Seed Size and Seeding Depth Influence on Canola Emergence and Performance in the Northern Great Plains. Agron J 96, 454–461. 10.2134/agronj2004.4540

24. Larroque, O.R., Calderini, D.F., Angus, J.F., 2022. Managing dryland wheat to produce high-quality grain. Field Crop Res 280, 108473. 10.1016/j.fcr.2022.108473

25. Li, N., Song, D., Peng, W., Zhan, J., Shi, J., Wang, X., Liu, G., Wang, H., 2019. Maternal control of seed weight in rapeseed (Brassica napus L.): the causal link between the size of pod (mother, source) and seed (offspring, sink). Plant Biotechnology Journal 17, 736–749. 10.1111/pbi.13011

26. López Pereira, M., Sadras, V.O., Trápani, N., 1999. Genetic improvement of sunflower in Argentina between 1930 and 1995. I. Yield and its components. Field Crop Res 62, 157–166. 10.1016/s0378-4290(99)00015-5

27. Massuia de Almeida, L.M., Avice, J.-C., Morvan-Bertrand, A., Wagner, M.-H., González-Centeno, M.R., Teissedre, P.-L., Bessoule, J.-J., Le Guédard, M., Kim, T.H., Mollier, A., Brunel-Muguet, S., 2021. High temperature patterns at the onset of seed maturation determine seed yield and quality in oilseed rape (Brassica napus L.) in relation to sulphur nutrition. Environ Exp Bot 185, 104400. 10.1016/j.envexpbot.2021.104400

28. McDonald, G.K., Genc, Y., Graham, R.D., 2008. A simple method to evaluate genetic variation in grain zinc concentration by correcting for differences in grain yield. Plant Soil 306, 49–55. 10.1007/s11104-008-9555-y

29. Mera, M., Lizana, X.C., Calderini, D.F., 2015. Chapter 6 - Cropping systems in environments with high yield potential of southern Chile. In: Sadras, V.O., Calderini, D.F. (Eds.), Crop Physiology (Second Edition). Academic Press, San Diego, pp. 111–140.

30. Merrill, A.L., Watt, B.K., 1973. Energy Value of Foods: Basis and Derivation. ARS United States Department of Agriculture, Washington DC.

31. Mińkowski, K., 2000. Influence of variety and size of winter rapeseed on content and chemical composition of hull and embryo. Rośliny Oleiste 21, 157–166.

32. Mondal, S., Pramanik, K., Panda, D., Dutta, D., Karmakar, S., Bose, B., 2022. Sulfur in Seeds: An Overview. Plants 11, 450. 10.3390/plants11030450

33. Musa Özcan, M., 2006. Determination of the mineral compositions of some selected oil-bearing seeds and kernels using Inductively Coupled Plasma Atomic Emission Spectrometry (ICP-AES). Grasas Aceites 57, 211–218. 10.3989/gya.2006.v57.i2.39

34. Newkirk, R., 2011. 8 - Meal Nutrient Composition. In: Daun, J.K., Eskin, N.A.M., Hickling, D. (Eds.), Canola. AOCS Press, pp. 229–244.

35. Panthee, D.R., Pantalone, V.R., West, D.R., Saxton, A.M., Sams, C.E., 2005. Quantitative Trait Loci for Seed Protein and Oil Concentration, and Seed Size in Soybean. Crop Sci 45, 2015–2022. 10.2135/cropsci2004.0720

36. Pokharel, M., Stamm, M., Hein, N.T., Jagadish, K.S., 2021. Heat stress affects floral morphology, silique set and seed quality in chamber and field grown winter canola. Journal of Agronomy and Crop Science 207, 465–480.

37. Quintero, A., Molero, G., Reynolds, M.P., Calderini, D.F., 2018. Trade-off between grain weight and grain number in wheat depends on GxE interaction: A case study of an elite CIMMYT panel (CIMCOG). Eur J Agron 92, 17–29. 10.1016/j.eja.2017.09.007

38. Rasheed, M.U., Lihong, W., Mahmood, A., Rehman, H.U., Javaid, M.M., Ali, B., Ishfaq, F., Ameen, M., Hassan, M.U., Hashem, A., Almutairi, K.F., Abd_Allah, E.F., 2024. Impact of Nitrogen, Sulphur, and Foliar Applied Thiourea on Growth, Oil Yield, and Fatty Acid Profile of Canola. Pol J Environ Stud. 10.15244/pjoes/186399

39. Richard, R., Lovegrove, A., Tosi, P., Casebow, R., Poole, M., Wingen, L.U., Griffiths, S., Shewry, P.R., 2025. Genetic analysis of grain protein content and deviation in wheat. J Cereal Sci 121, 104099. 10.1016/j.jcs.2024.104099

40. Richards, R.A., Lukacs, Z., 2002. Seedling vigour in wheat - sources of variation for genetic and agronomic improvement. Aust J Agr Res 53, 41–50. 10.1071/AR00147

41. Rivelli, G., Calderini, D.F., Abeledo, L.G., Miralles, D.J., Rondanini, D.P., 2024. Yield and quality traits of wheat and rapeseed in response to source-sink ratio and heat stress in post-flowering. Eur J Agron 152, 127028. 10.1016/j.eja.2023.127028

42. Rivelli, G.M., Fernández Long, M.E., Abeledo, L.G., Calderini, D.F., Miralles, D.J., Rondanini, D.P., 2021. Assessment of heat stress and cloudiness probabilities in post-flowering of spring wheat and canola in the Southern Cone of South America. Theoretical and Applied Climatology 145, 1485–1502. 10.1007/s00704-021-03694-x

43. Rondanini, D.P., del Pilar Vilarino, M., Roberts, M.E., Polosa, M.A., Botto, J.F., 2014. Physiological responses of spring rapeseed (Brassica napus) to red/far-red ratios and irradiance during pre- and post-flowering stages. Physiol Plantarum 152, 784–794. 10.1111/ppl.12227

44. Rotkiewicz, D., Tańska, M., Konopka, I., 2002. Seed size of rapeseed as a factor determining their technological value and quality of oil. Rośliny Oleiste 23, 103–112.

45. Sadzawka, A., Carrasco, M.A., Demanet, R., Flores, H., Grez, R., Mora, M.d.l.L., Neaman, A., 2007. Métodos de análisis de tejidos vegetales. Instituto de Investigaciones Agropecuarias. Centro Regional de Investigación La Platina, Santiago, Chile.

46. Sandaña, P.A., Harcha, C.I., Calderini, D.F., 2009. Sensitivity of yield and grain nitrogen concentration of wheat, lupin and pea to source reduction during grain filling. A comparative survey under high yielding conditions. Field Crop Res 114, 233–243. 10.1016/j.fcr.2009.08.003

47. Thiyam-Hollaender, U., Eskin, N.A.M., Matthäus, B., 2013. Canola and rapeseed : production, processing, food quality, and nutrition. CRC Press, Boca Raton.

48. Triboi-Blondel, A.-M., Renard, M., 1999. Effects of temperature and water stress on fatty acid composition of rapeseed oil.

49. Vahedifar, A., Wu, J., 2022. Chapter Two - Extraction, nutrition, functionality and commercial applications of canola proteins as an underutilized plant protein source for human nutrition. In: Wu, J. (Ed.), Advances in Food and Nutrition Research. Academic Press, pp. 17–69.

50. Verdejo, J., Calderini, D.F., 2020. Plasticity of seed weight in winter and spring rapeseed is higher in a narrow but different window after flowering. Field Crop Res 250, 107777. 10.1016/j.fcr.2020.107777

51. Verdejo, J.F., Calderini, D.F., 2025. Resilience of rapeseed to temperature increase during early grain filling in a high yielding environment. Field Crop Res 330, 109950. 10.1016/j.fcr.2025.109950

52. Weiss, E., 2000. Rapeseed. Oilseed Crops. Blackwell Science Ltd., Victoria, Australia.

53. Zhang, W.-H., Zhou, Y., Dibley, K.E., Tyerman, S.D., Furbank, R.T., Patrick, J.W., 2007. Nutrient loading of developing seeds. Funct Plant Biol 34, 314–331. 10.1071/FP06271

54. Zhu, L., Zhang, D., Fu, T., Shen, J., Wen, J., 2011. Analysis of yield and disease resistance traits of new winter rapeseed varieties over the past twenty years in China. Agricultural Science & Technology Hunan 12, 842–846.

